# CycleMix: Gaussian Mixture Modeling of the Cell Cycle

**DOI:** 10.1101/2025.02.11.637734

**Authors:** Jack Edward Peplinski, Tallulah Andrews

**Affiliations:** Department of Electrical and Computer Engineering & Ivey School of Business, University of Western Ontario; Department of Biochemistry & Department of Oncology, Schulich School of Medicine and Dentistry, University of Western Ontario; Department of Computer Science, University of Western Ontario

## Abstract

The cell cycle is a crucial component of many biological processes, including cancer, tissue repair, and inflammation. However, due to the heterogeneity of this cycle it has been difficult to assess the extent of proliferation in clinical tissues. Single-cell RNAseq (scRNAseq) and spatial transcriptomics enables high resolution measurement of gene expression enabling the classification of individual cells into their cycling state. The most widely used method for cell-cycle assignment is Seurat, however their approach is limited to classifying cells into only 3 states: G1, S, G2M and frequently incorrectly labels fully mature non-cycling cells (e.g. neurons) as S/G2M. Here we propose CycleMix, an alternative cell-cycle assignment algorithm that can flexibly assign cells into any number of states provided sufficient marker genes as well as being capable of identifying when cells are not cycling.

## Introduction

The cell cycle is a fundamental biological process that enables the survival and reproduction of all living things. Dysregulation of the cell cycle has been implicated in diseases such as cancer [1,2], neurodegeneration [3,4], polycystic kidney disease [5], and schizophrenia [6]. More recently, cell cycle length has been hypothesized to control cell differentiation [7]. However the cell cycle can also be an unwanted confounding variable when working with cells in culture or in highly proliferative tissues [8–10].

Single-cell RNA sequencing (scRNAseq) enables high-throughput measurement of heterogeneity within individual cells [11–15]. Datasets now routinely include upwards of 100,000 single cells obtained from dozens of individual samples [16–20]. The cell cycle represents a major source of variability within these datasets, both as variability between proliferative cells in different points in the cell cycle [21,22] as well as variability between proliferative and non-proliferative cells [23]. Most cells in healthy adult tissues are non-proliferative, whereas a relatively small proportion of stem and progenitors actively proliferate to maintain tissue homeostasis.

Activation of proliferation can be a crucial step in normal and diseased biological processes, for instance T-helper cells have been observed to increase proliferation at the point of cell-fate decision [24]. In cancer, the activation, dysregulation and suppression of proliferation affects drug persistence [25], progression and invasion [26], and successful metastasis [1]. Thus the cell cycle was one of the first biological systems targeted with single-cell analysis using synchronized embryonic stem cells [27,28], fluorescent labeling of key cell cycle markers [29], and flow cytometry properties [30].

In parallel, multiple statistical approaches have been developed to classify single cells into their cell-cycle phase, which can be divided into two main approaches: continuous cycle inference and discrete cell classification. Oscope [29] identifies oscillating genes using a two-dimensional sine-wave model which are then used to order cells along the cycle. DeepCycle [31] combined inferred gene expression dynamics from RNA velocity with angular pseudotime to infer an ordering of cells through the cell cycle. Both are examples of continuous cycle inference methods, which require prior knowledge of which cells are actively cycling and which are quiescent. In contrast, classification based methods such as cyclone [32], which uses relative expression of gene pairs, and Seurat [33], which uses average normalized expression of marker genes, assign to each cell a specific phase: G1, S, G2/M. These methods do not explicitly allow cells to be classified as non-cycling, but typically cells in the “G1” phase are assumed to primarily represent non-cycling cells, rather than specifically proliferating cells within the G1 phase. This is particularly true of Seurat which does not use any markers to define the G1 phase but simply classifies all non-S and non-G2M cells as G1.

We present a novel cell-cycle classification method, CycleMix, which uses mixture models to determine which cells in a single-cell experiment are actively cycling and which stage those cells are currently in. We assess the performance of CycleMix on low-throughput scRNAseq datasets where cell-cycle phase was identified using orthogonal experimental approaches, and on high-throughput scRNAseq datasets where a priori biological knowledge and author annotations can be used to identify proliferative and quiescent cell-types. We show that CycleMix is highly scalable, performs competitively on low-throughput datasets and is the best performing method on high-throughput datasets compared to current state-of-the-art methods.

## Results

### CycleMix

CycleMix classifies cells into cell-cycle phases using a slight variation on cell-cycle scoring used by others [34–36]. Raw UMI or read counts are library-size normalized to the median library size across cells, then log transformed (**Figure 1**). Phase scores are calculated using a weighted mean enabling both up and downregulated markers for each phase. Marker genes lists from multiple sources are available within CycleMix including those used by Seurat, Macosko et al. (2015) [35], Tirosh et al. (2016) [34], and defined by Whitfield et al. (2002) [37]. As noted by others, we find the Tirosh dataset works well and use it by default in this manuscript [38]. Whitfield contains signatures for more fine-scale dissection of the cell cycle however these are often less cleanly separable in noisy single-cell data.

**Figure 1:**
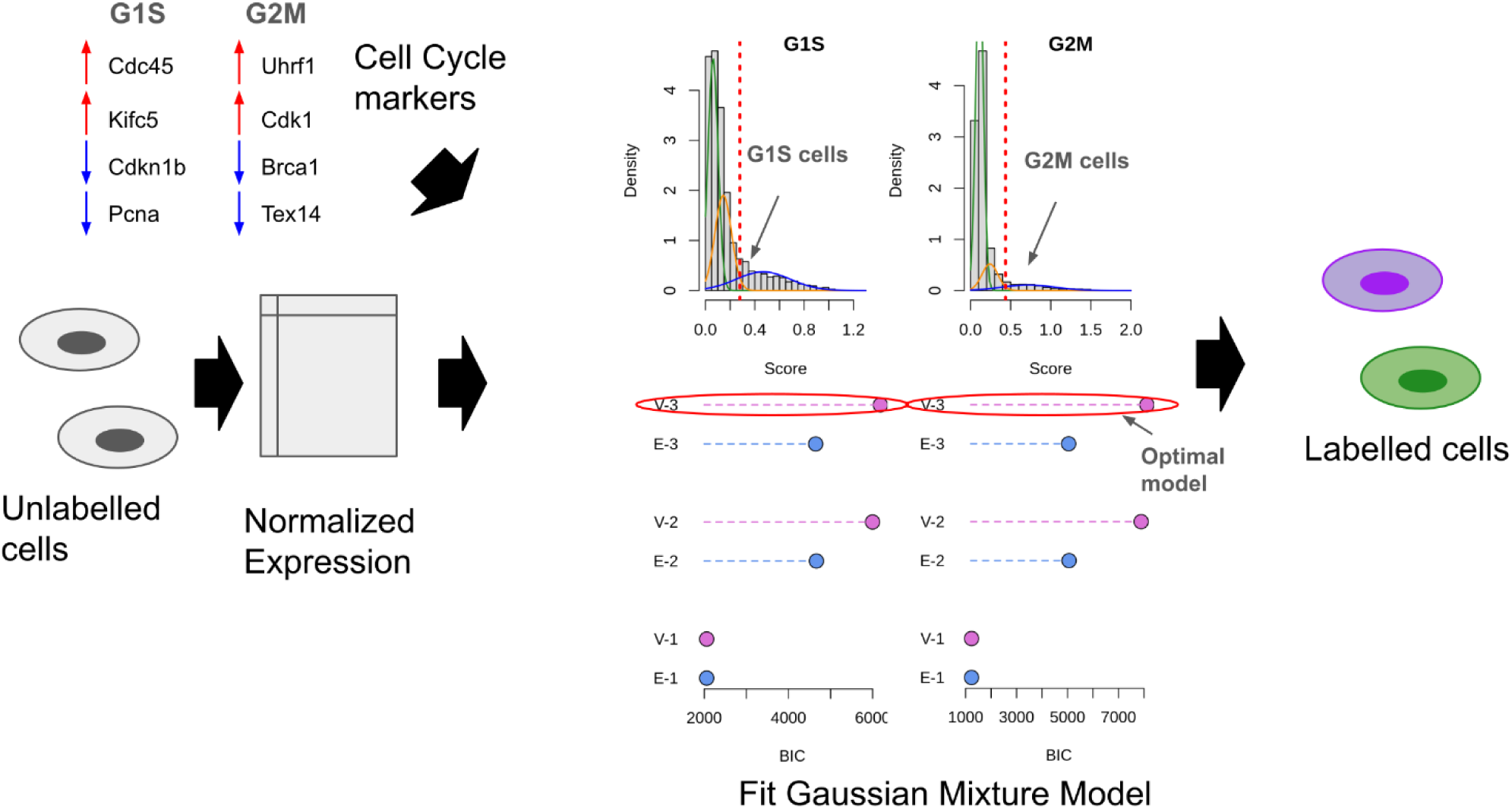
The CycleMix approach to cell-cycle classification. Log2-normalized expression is calculated for each cell, optionally any other normalization such as batch effect correction are applied. The weighted average of the normalized expression across up and down regulated markers of each cell-cycle phase. Mixture models of 1, 2, or 3 Gaussian distributions are fit to the average scores across cells. The Bayesian Information Criterion (BIC) determines the optimal model to describe the data. Cells assigned to the Gaussian distribution with the highest mean for a phase are assigned to that phase.

Scores of each phase are then fit to six different Gaussian mixture models corresponding to mixtures of 1, 2, or 3 Gaussian distributions with either equal or variable variance. The Bayesian Information Criterion (BIC) which incorporates both the likelihood of the data given the model and the number of parameters used to fit the model to the data is used to select the optimal model. Typically, the three Gaussian mixture variable variance model is chosen with distributions corresponding to cells in the specific phase with the highest average scores, cells that are cycling but not in this specific phase in the middle distribution and non-cycling cells with the lowest mean score (**Figure S1**). Cells in the distribution with the highest mean are assigned to that phase. Phase scores are then Z-transformed and cells assigned to multiple phases are assigned the phase with the highest Z-score.

Gaussian or negative binomial regression models can then be used to remove differences between any set of cell-cycle phases the user would like, enabling removal of cell-cycle phase differences while preserving differences between cycling and non-cycling cells.

### Benchmarking on datasets with know cell-cycle phase

To validate the effectiveness of CycleMix compared to other cell-cycle inference methods, we obtained five publicly available datasets where cell-cycle stage was assessed using an orthogonal method by synchronizing cells, flow sorting using cell shape features, or using fluorescent reporter genes (**Table 1**). Each dataset was classified into cell-cycle phases using CycleMix, Seurat [36], and cyclone [32]. Overall, cyclone had the highest accuracy in classifying single-cells across all datasets (**Figure 2A**), with CycleMix a close second, whereas Seurat performed no better than random chance on both Buettner and Chen datasets. While cyclone was most accurate it was also much less scalable than CycleMix or Seurat (Figure 2B) requiring four minutes to classify the 332 cells of the Chen dataset.

**Table 1:** Benchmarking Datasets.

| Dataset | Description | Database Accession |
| --- | --- | --- |
| Sasagawa [27] | 56 synchronized mouse ES cells by Quartz-seq | GSE42268 (GEO) |
| Chen [39] | Synchronized mouse spermatocytes by Smartseq2 | GSE107644 (GEO) |
| Buettner [28] | Synchronized mouse ES cells by Fluidigm C1 | E-MTAB-2805 (ArrayExpress) |
| McDavid [30] | Flow sorted human cell-lines by Nanostring nCounter targeted scRNAseq | https://doi.org/10.1371/journal.pcbi.1003696.s009 |
| Leng [29] | FUCCI H1 hESCs by Fluidigm C1 | GSE64016 (GEO) |

**Figure 2:**
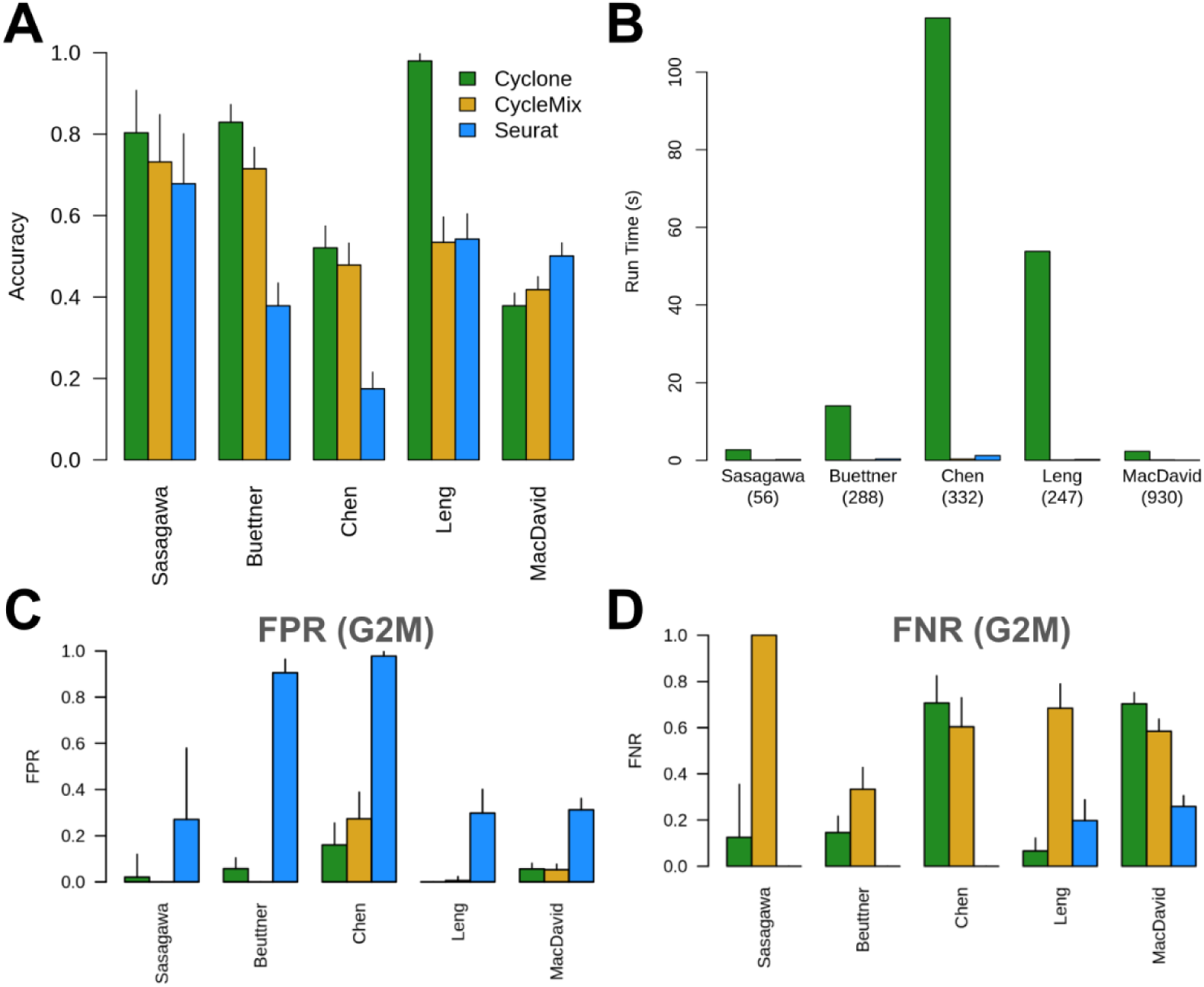
Benchmarking on 5 datasets with orthogonally determined cell-cycle phase. (A) Overall accuracy of cell-cycle stage predictions. (B) Runtime of each method on each datasets. (C) Relative number of G1 or S cells assigned to G2M (D) Proportion of G2M cells assigned to G1 or S. Green = Cyclone, Gold = CycleMix, Blue = Seurat. Bars indicate 95% CI.

Seurat consistently overestimates the proportion of mitotic cells (**Figure 2C**) assigning 35%-98% of cells to the G2M stage across the datasets. In contrast, CycleMix was more conservative, and tended to underestimate the proportion of mitotic cells (**Figure 2D**). cyclone was inconsistent in it’s errors, overestimating the number of S-stage cells in the Leng and Sasagawa, but underestimating both S and G2M in Chen and MacDavid datasets.

### Downsampling datasets to mimic droplet-based technologies

While the above datasets have reliable ground-truth cell-cycle stage information available the cells were captured and sequenced using plate-based [27,39] and microfluidic chip based methods [28,29] without the use of unique molecular identifiers. As a consequence, these datasets are much smaller (56-930 cells), have different noise profiles and sensitivities than current state-of-the-art droplet-based methods. We repeatedly downsampled three of these datasets to generate data more similar to those from droplet-based methods (**Figure S2**). Results were similar to those observed for the original datasets with CycleMix and cyclone performing well on the Buettner datasets, none performing well on the Chen dataset and cyclone outperforming the others on the Leng dataset.

### Benchmarking on droplet-based scRNAseq datasets

To benchmark the cell-cycle classification methods on state-of-the-art scRNAseq datasets we collected six publicly available datasets containing cell-types known to be highly proliferative and mature quiescent cell-types (**Table 2**). We ran each cell-cycle classification method on all cells in each dataset and assessed the proportion of cells assigned to S/G2M phases among the mature non-proliferative cell-types and the expected highly proliferative cell-types (**Figure 3A**). We found that both CycleMix and Seurat but not cyclone consistently assigned a higher proportion of cells to S/G2M for the expected proliferative cell-types than the quiescent cell-types. This suggests that cyclone may suffer from being trained on full-length Smartseq2 scRNAseq and does not perform well on 3’-tag based scRNAseq datasets. However, only CycleMix accurately assigned more than >90% of quiescent cells to a non-cycling phase, whereas Seurat assigned 25%-50% of quiescent cells to S/G2M.

**Table 2:**
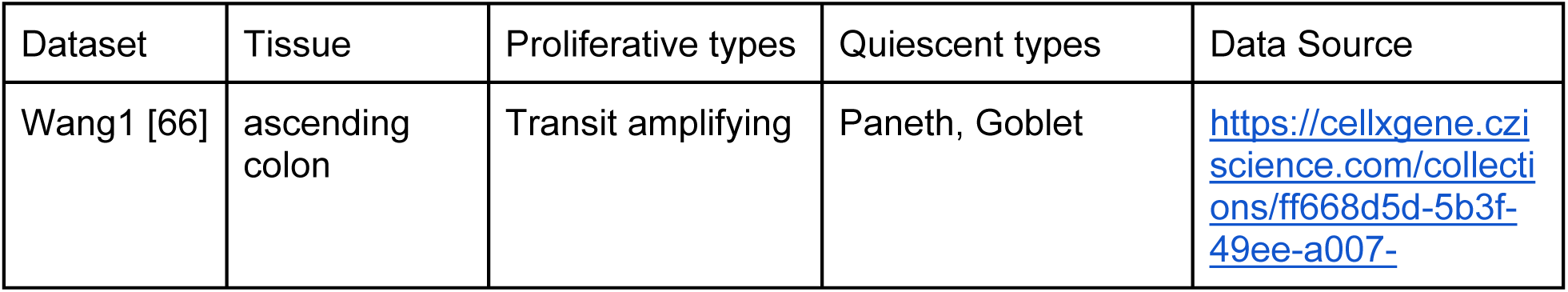
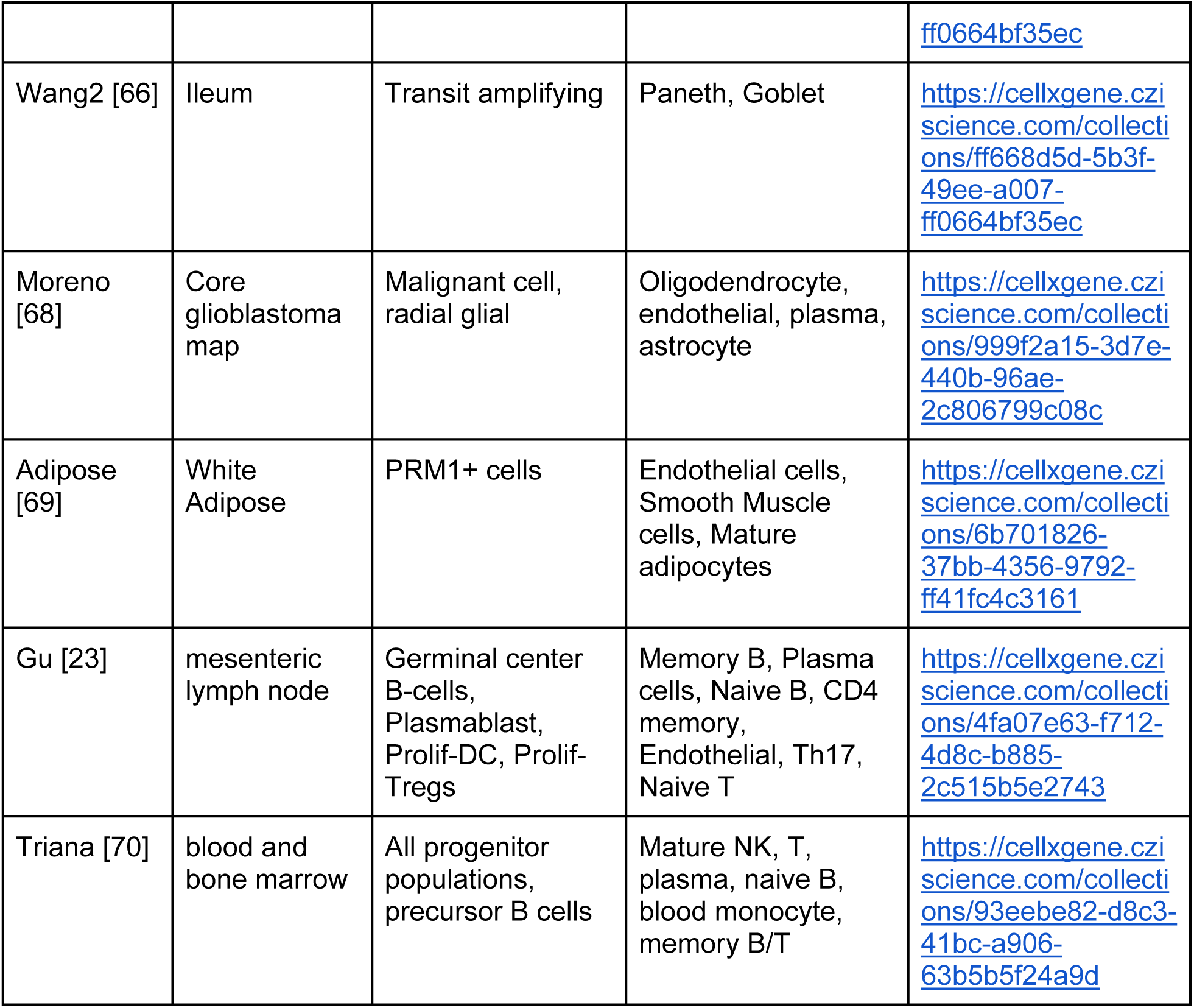
Real datasets containing populations of proliferating cells.

**Figure 3:**
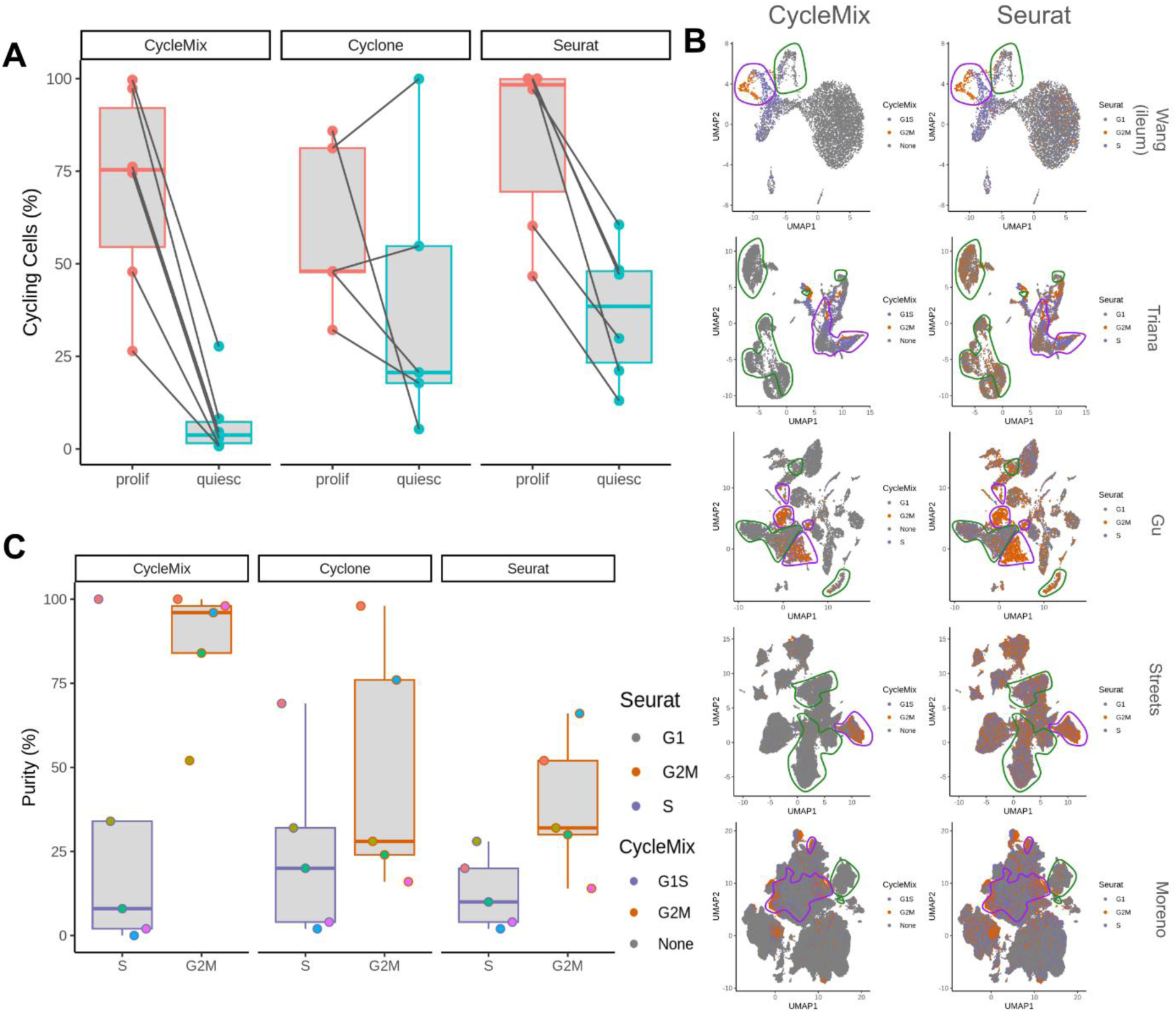
Assessing performance on droplet-based datasets with known proliferative and quiescent cell populations. (A) Proportion of cells classified as S or G2M by each method that were annotated as belonging to a proliferative or quiescent cell-type. Lines connect matching points for each dataset. (B) UMAPs depicting cell-cycle classification by CycleMix (left) and Seurat (right) for the different datasets. Purple and green outlines indicate the proliferative and quiescent populations respectively. (C) Local purity of cell-type classifications.

Since even among highly proliferative cell-types it is unlikely that every cell was currently actively cycling when the scRNAseq was performed, we also examined the local neighbourhoods of S and G2M cells (**Figure 3B-C**). Since the cell-cycle involves consistent changes in protein expression to perform DNA replication and then mitotic division, we reasoned that cells of the same cell-type within the same stage of the cell-cycle should cluster together in gene-expression space. Thus we expect the local neighbourhood around G2M cells to predominantly contain other G2M cells and likewise for cells currently in S phase. For all methods, we observe that G2M cells were significantly more likely to cluster together than S-phase cells (p < 10^−10^, Wilcox-rank-sum test). This was most pronounced using the classifications obtained from CycleMix, where on average >50% of neighbouring G2M cells were other G2M cells across all datasets, in contrast to ∼37% for cyclone and ∼25% for Seurat. These results were consistent with Seurat exhibiting much higher false-positive rates for S and G2M cell-type assignments as we observed in the gold-standard datasets.

### Cell-cycle Regression

The best method to account for the cell cycle in scRNAseq data remains uncertain [12,13]. A common approach is to generate a quantitative score representing the similarity of the cell to each cell-cycle stage then use regression models to remove the effect of these quantitative scores from the expression of each gene (e.g., Seurat [36]). Alternatively, discrete classifications can be used for this regression. Other approaches have included factor decomposition followed by removal of cell-cycle related factors [28]. A key challenge with these methods is when the cell cycle is confounded with biological variation of interest. For instance highly proliferative subpopulations of stem cells or reactive immune cells, as regression models may not distinguish the problematic cell-cycle variability from the biologically relevant cell-type variability. Within our CycleMix package, we include a partial regression model based on discrete classifications which only removes differences between S and G2M cell-cycle phases while preserving differences between cycling and non-cycling cells (see Methods).

Comparing our discrete partial regression to the Seurat score-based full regression, we find CycleMix is more effective at intermingling G2M and S cells while preserving the integrity of highly proliferative cell-types (**Figure 4**). Both regression methods significantly increased the mixing of cells in different stages of the cell cycle (Seurat: p = 0.02, CycleMix: p = 0.01, Figure 4B). Whereas only CycleMix preserved, and sometimes enhanced, the consistency of proliferative populations (**Figure 4A, C**).

**Figure 4:**
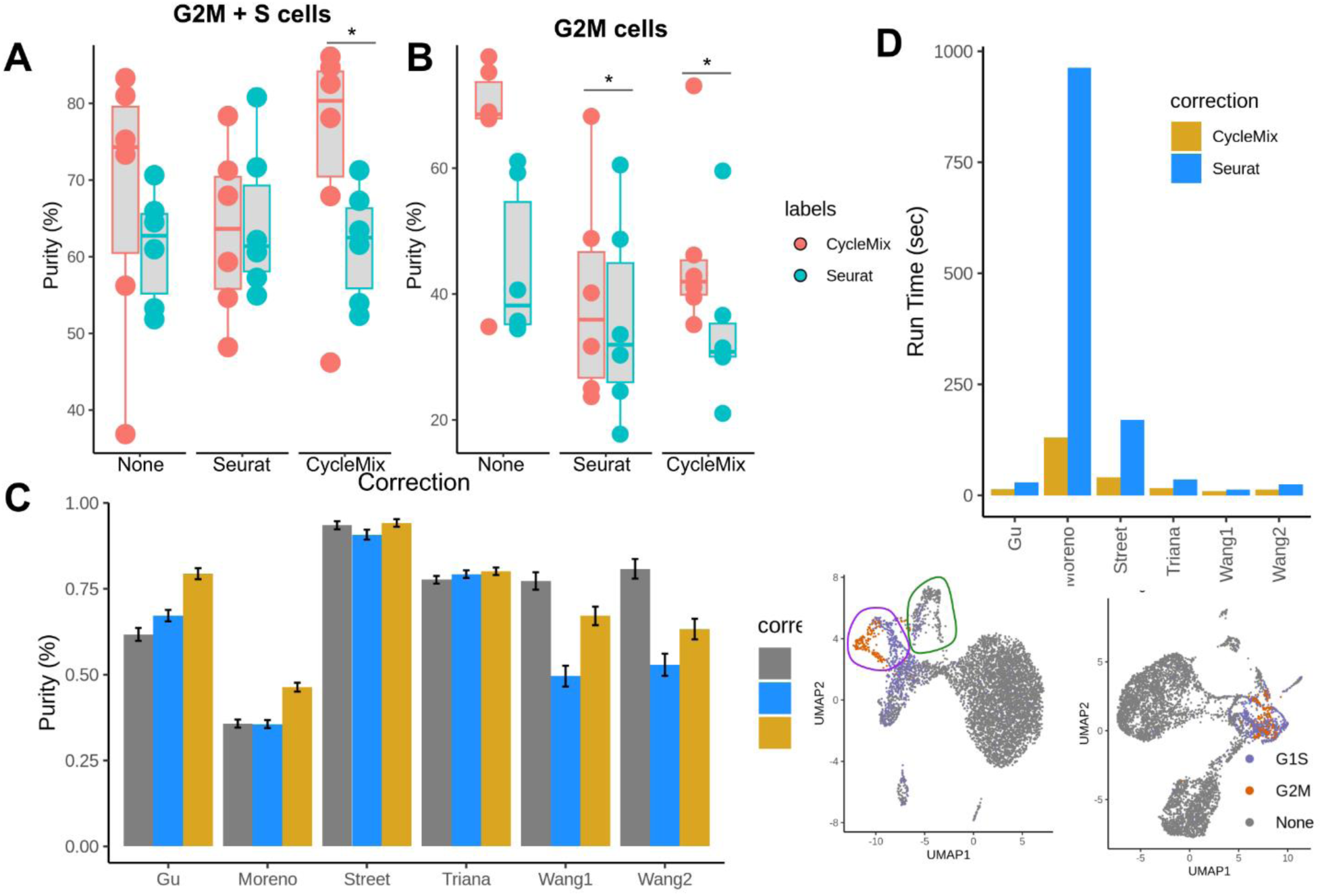
Correcting for cell cycle differences. (A) CycleMix regression preserves separation of G2M/S cells from non-proliferating cells. Each dot represents one dataset, colours indicate the origin of the cell-cycle labels used to determine G2M/S cells, None = uncorrected, Seurat = Seurat CC regression. CycleMix = partial CC regression included in CycleMix. Stars indicate significant differences compared to no-regression (p < 0.05, paired t-test) (B) As in A but considering only purity of G2M cells within themselves. (C) CycleMix regression preserves or enhances separation of proliferative cell-types. Average purity of each proliferative cell-type. Error bars indicate 95% confidence intervals. (D) Runtime for CycleMix and Seurat regression on each dataset.

### Identifying Quiescent Cells

CycleMix can be used to assign cells to any category defined by a set of up and/or down regulated genes. Using a recent publication that identified genes consistently associated with quiescence in 3 different stem cell populations [40], we were able to use CycleMix to identify quiescent (G0) cells in each of our six droplet-based datasets (**Figure 5A**). Cell-types expected to be quiescent were enriched in these cells vs. proliferative cell-types as expected (**Figure 5A**). These markers were defined based stem cells thus may not be ideal for assessing quiescence in differentiated cells. We used MAST [41] to identify genes associated with quiescence after controlling for cell-type and sample-specific effects in all our datasets. Only AHNAK, B2M, MALAT1 and NEAT1 were consistently upregulated in quiescent cells (**Figure 5B**). MALAT1 and NEAT1 are nucleus-specific transcripts which are associated with cell-integrity in scRNAseq experiments thus are unlikely to specifically identify G0 cells [42]. In contrast, 152 genes were consistently downregulated in quiescent cells (**Table S1**), as expected 32% of these genes are known targets of Myc, a well known oncogene and pro-proliferation transcription factor [43–45], and 29% were targets of the proliferation-associated E2F transcription factor [46] (**Figure 5C**) suggesting that in differentiated cells quiescence is mainly determined by down regulation of proliferation pathways.

**Figure 5:**
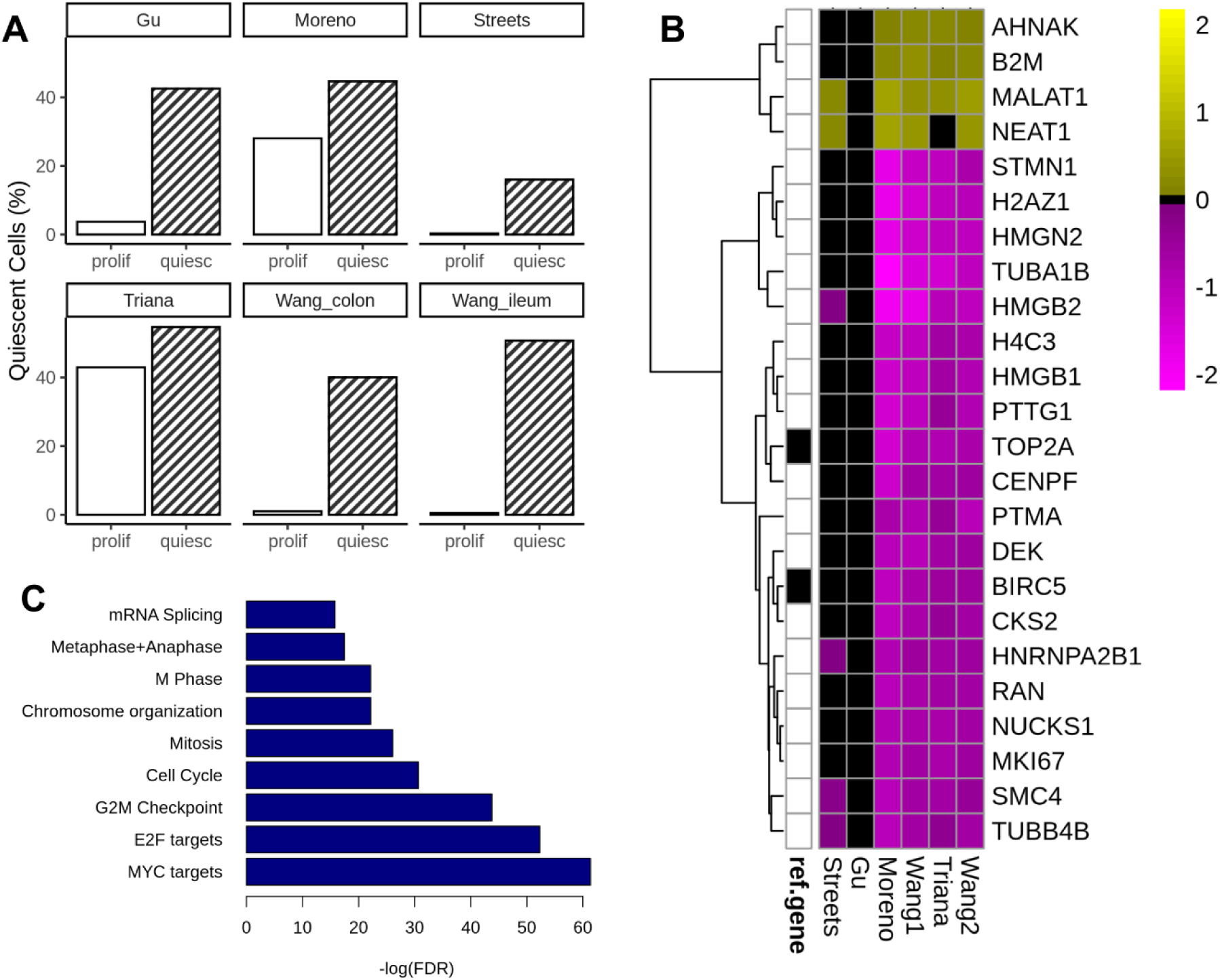
CycleMix can identify G0 cells. (A) Proportion of cells labelled as quiescent among cell-types that should be proliferative or quiescent in each dataset. Quiescent genes were obtained from Cheug and Rando (2013). (B) Top most consistently differentially expressed genes between quiescent and proliferative cells across datasets. (C) Top significantly enriched pathways among genes downregulated in quiescent cells, ref.gene indicates whether it was among the genes identified in Cheug and Rando.

### Proliferation of iPSC-derived Kupffer cells

Kupffer cells are unique liver-resident macrophages that phagocytose worn out and damaged red blood cells in the liver. Kupffer cells are embryonically derived and locally self-renew to maintain their population in the liver independent of the bonemarrow [47,48]. However the mechanisms and frequency of this self-renewing proliferation is not known. Recently, we generated scRNAseq data of iPSC-derived Kupffer cells that were transplanted into a humanized mouse [49], these cells expressed gene signatures consistent with fully mature liver Kupffer cells and could phagocytose red blood cells. As these cells were colonizing the host liver, we expect to observe some portion of proliferating cells among this population. Using CycleMix, we can identify a distinct subpopulation of actively proliferating cells accounting for 3.5% of the Kupffer cell population, whereas Seurat classifies 56% of the Kupffer cells as actively proliferating (**Figure 6A,B**). Expression of the top positive and negative markers of quiescence identified above confirm predominantly quiescent signature for most cells and only a small population of proliferating cells (**Figure 6C,D**). Comparing to normal human development [50], revealed that using CycleMix iPSC-derived Kupffer cells were significantly less proliferative than normal human fetal Kupffer cells (p < 0.05), whereas using Seurat would lead to the opposite conclusion (**Figure S4**).

**Figure 6:**
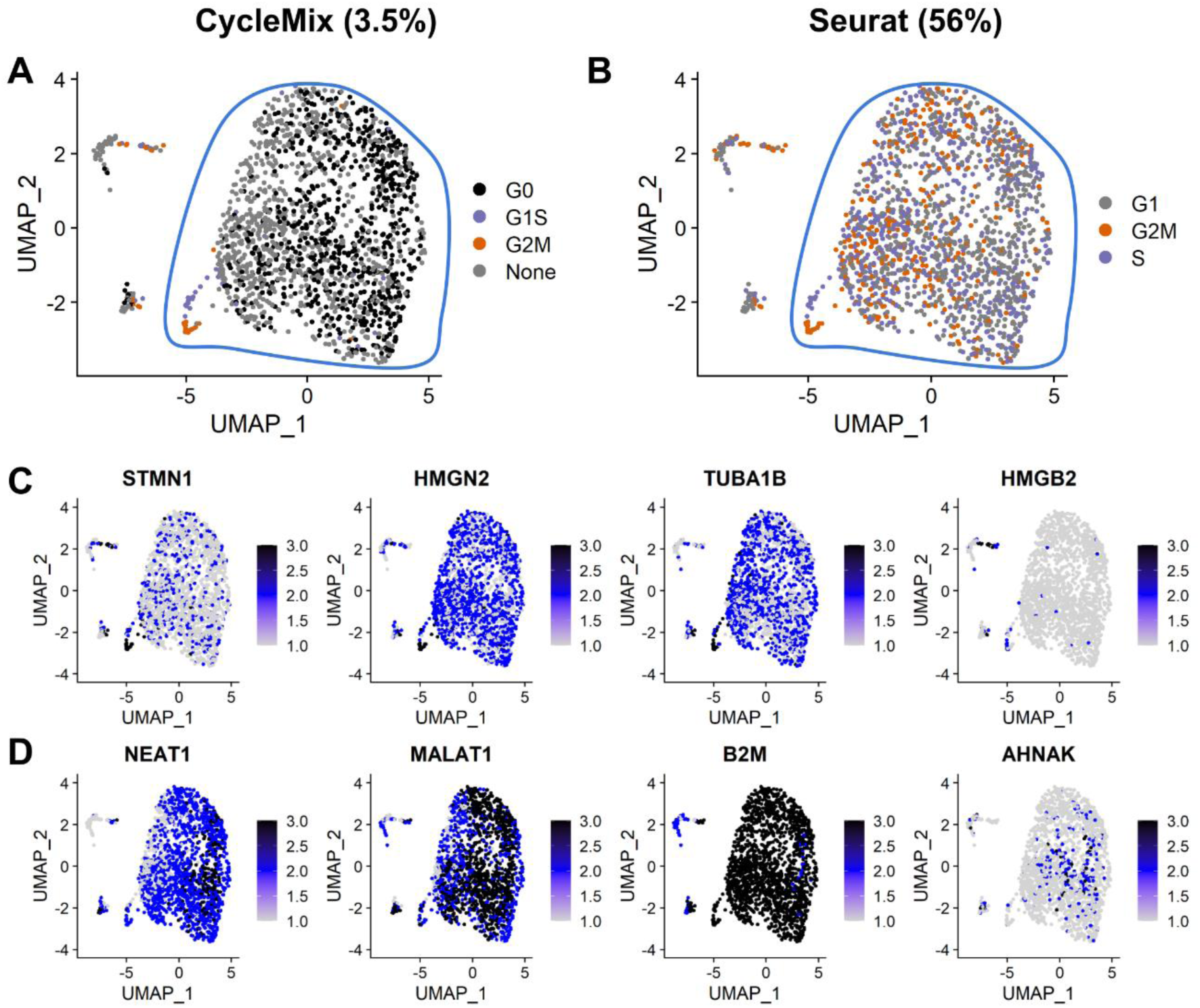
CycleMix identified subpopulation of proliferating iPSC-derived Kupffer cells. (A) Cell cycle classification of iPSC-derived Kupffer cells (blue circle) using CycleMix. (B) As in A but with Seurat classifications. (C) Expression of top 4 downregulated quiescence genes (D) Expression of top 4 up regulated quiescence genes.

## Discussion

We present CycleMix, a novel scalable cell-cycle classification algorithm based on Gaussian Mixture modeling. Briefly, this approach uses a weighted average log-normalized expression to combine positive and negative gene markers to generate stage-specific scores which are binarized into discrete classifications by fitting a mixture of Gaussian distributions and using the BIC to select the optimal model (**Figure 1**). This enabled it to more accurately distinguish cycling vs. non-cycling cells than the most commonly used approaches, while maintaining a high degree of scalability. We demonstrated its performance on both gold and silver standard datasets from a range of different single-cell RNAseq technologies.

Of note, we observed that cyclone outperformed CycleMix on most gold-standard benchmarking datasets (**Figure 2**), but performed poorly on silver-standard datasets (Figure 4). This is due to all current gold-standard datasets using full-length transcript sequencing methods that do not employ unique molecular identifiers. This results in significantly average expression levels and noise profiles compared to droplet-based scRNAseq approaches with full-length data best described by a zero-inflated negative binomial and umi-tagged data not exhibiting zero-inflation [51–53]. cyclone was trained exclusively on full-length Smartseq2 scRNAseq datasets, to identify gene pairs where the relative expression between the genes changes between cell-cycle phases [32]. Thus, it is not surprising that it fails to generalize to UMI-tagged data.

Current state-of-the-art scRNAseq uses almost exclusively 3’ or 5’ tag-based scRNAseq datasets, when we tested all three classification methods on such data CycleMix clearly outperformed the others. In contrast, the most widely used method implemented in Seurat demonstrated very high false-positive rates with 40% of quiescent cells being classified as cycling, in contrast to <10% for CycleMix (**Figure 2**). This was consistent with results on the full-length gold-standard datasets where Seurat classified an average of 55% non-G2M cells as G2M, vs. 6% for CycleMix and 5% for cyclone (**Figure 2**). This is likely a result of the flat threshold of an average score of 1 for either S or G2M to classify a cell as cycling.

It has been noted that there are two different forms of the cell-cycle in mammalian cells: a fast version with minimal G1 phase used by embryonic stem cells (ESCs) and cancer, and one with a significant G1 used by normal adult tissues [38,54,55]. This fast cell-cycle may directly contribute to maintaining pluripotency by blocking the transcription of long genes associated with differentiation [7]. Three of the five gold standard datasets originate from ESCs or induced pluripotent cells, and a fourth used cancer-derived cell-lines in contrast our silver-standard datasets were predominantly healthy adult tissues with only a single cancer dataset. The gene-list we used for cell-cycle classification was derived from Tirosh (2016) [56] which examines metastatic cancer, thus likely represent the fast version of the cell-cycle which may explain why the G2M classifications were more consistent and concentrated within the datasets than the G1S phase cells. Cell-cycle phase markers derived from cycline normal adult tissues may further improve classification accuracy.

Using a recently published set of quiescence markers [40], CycleMix could classify cells across all tissues as G0 vs. G1S or G2M (**Figure 3**), however, these classifications were primarily driven by down regulator of various pro-proliferation markers rather than any consistent positive markers of G0 across tissues. Indeed, the only transcripts consistently associated with G0 were two nuclear-associated long-noncoding RNAs and AHNAK and B2M. AHNAK has been previously associated with anti-proliferative effects in cancer [57,58], but it’s unclear if it has a direct interaction with cellular proliferation machinery [59]. B2M is a member of the major histocompatibility complex (MHC-I), that may have indirect effects on suppressing proliferation [60]. Thus, it is unclear whether there are any consistently upregulated genes associated with quiescence across all tissues and cell-types.

CycleMix can also be used to annotate cells to cell-type, if provided with an appropriate list of marker genes (**Figure S5**); however, accuracy on the datasets used here was not very high due to the high frequency of overlapping marker genes between similar cell-types. This resulted in many cells being misclassified into more general cell-types as markers defining fine-grained cell subtypes are fewer in number and often more lowly expressed than major cell-lineage defining markers and is a challenge for all automatic cell-annotation methods [61].

Thus, we have created a more accurate and more flexible single cell-cycle classification approach compared to the current state of the art. We demonstrated the importance of accurate cell-cycle classification when considering determining the maturity of iPSC-derived cells. Other instances where accurate absolute quantification of proliferation is important is in cancer biology, due to known correlations between high tumour proliferation and poor patient prognosis [62,63], and potential importance for inflammation-associated disease as proliferation of lymphocytes and subsequent suppression is a key step in the immune response [64]. We have shown that only CycleMix can accurately estimate the absolute frequency of proliferation within scRNAseq datasets.

## Methods

### Classifying Cells

Gene signatures for each phase were obtained from the published literature, including Whitfield et al. [37], Tirosh et al. [56], Macosko et al. [35] and from the Seurat(v4) package [36]. Quiescence/G0 signatures were obtained from Reactome’s “G0” gene list, and Cheung et al. [65]. Genes upregulated in the specific phase were assigned a weight of +1, while genes downregulated in a specific phase were assigned a weight of −1.

Normalized gene expression values are extracted from a provided SingleCellExperiment or Seurat object, and the weighted average expression across genes is calculated for each phase for each cell. For each phase, a Gaussian mixture model is fit to the resulting cell-specific scores (relying on the central limit theorem). We consider mixtures of 1, 2, or 3 Gaussians and the optimal model is identified using the Bayesian Information Criterion (BIC). Since any population of cycling cells will have cells in at least two phases: S & G2M, if the BIC identifies a single Gaussian as the best fit model then no cells are assigned to that phase. Otherwise, cells assigned to the Gaussian with the highest mean are assigned to this phase.

As this approach can lead to a single cell being assigned to multiple phases, the user has the choice to keep all assigned phases or as we do in this manuscript assign cells to the phase with the highest score after rescaling each set of scores. If a cell is not assigned to any phase it is labeled as “None”.

### Regressing Cell-Cycle Phase

CycleMix provides the option to regress out one or more phases of the cell cycle, either using discrete classifications or the quantitative scores for each phase. For discrete classifications, one phase is chosen as the reference stage, and differences between this stage and each other specified stage are calculated using negative binomial regression and this difference is removed. Alternatively, cell-cycle differences can be removed by regressing out the scores assigned to each cell for each cell-cycle phase. This is calculated using negative binomial regression for the specified phases. In both cases, the regression coefficients are adjusted to account for the number of 0 present in the raw gene expression data, negative expression is not biologically meaningful, as below:

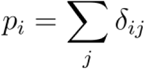

Where *δ_ij_* indicates whether the expression of gene i in cell j is greater than 0.

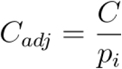

When regressing cell-cycle scores, we replace *C* above with the residual:

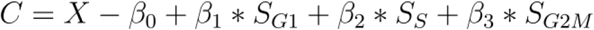

### Benchmarking

#### Datasets

Publicly available datasets with orthogonally determined cell-cycle phase were obtained from appropriate databases (**Table 1**).

#### Timing & Scalability

All methods were run on a HPC cluster with 1 CPU (Intel Gold 6148 Skylake 2.4 GHz) and 16 Gb of RAM. Algorithm timing included only the cell-cycle phase assignment, all data preprocessing steps were excluded. To test scalability, the Buettner dataset was resampled as described below generating increasing numbers of cells.

#### Resampling Data

For Chen, Buettner, and Leng datasets we generated synthetic data more similar to current state-of-the-art droplet-based scRNAseq. We obtained estimates of total UMIs/cell from 10,000 cells from two recent scRNAseq datasets generated using the 10X Chromium platform of human intestine [66] and mouse lymph node [23] (**Table 2**). We randomly sampled from these values for 1,000 cells of each cell-cycle phase (G1, S, G2M) for each dataset. For each synthetic cell we randomly selected one cell of the appropriate stage from the respective ground truth cell-cycle dataset and randomly sampled the specified number of reads from the read counts of the selected cell weighting each read equally, thus each gene is weighted by its total number of reads. This resulted in a similar number of genes detected per cell as in the original 10X datasets (**Figure S1**). Each cell-cycle classification method was run on the resampled datasets and performance was evaluated as above. The MacDavid dataset was excluded since it did not measure the whole transcriptome, and the Sasagawa dataset was excluded due to the low number of G2M and S samples (<10 samples for each).

#### Real datasets

Annotated droplet-based single-cell datasets (**Table 2**) were downloaded by cellxgene [67]. For Cyclemix, datasets were normalized to the median total UMI counts across cells, the log2 transformed with a pseudocount of 1. For Seurat, we used the provided log2-normalized gene expression from cellxgene. cyclone was run on raw counts. All methods were run on the complete dataset for the specified tissues. Cell-types were classified as proliferative or quiescent or unknown based on known biology and confirmed by inspecting the expression of TOP2A and HGMB2. cyclone was not run on the Moreno dataset as this dataset contained > 100,000 cells, making it computationally prohibitive to run cyclone on it.

Local purity of cell-cycle phases were calculated to account for cell-type such as malignant tumour cells where it is expected that only a subset of cells would be actively cycling. K-nearest neighbours of each cell was calculated using Seurat with a k=100. For each cell of the appropriate phase, we calculate what proportion of their neighbours were assigned to the same phase. As the datasets contained over 4,000 cells k=100 represents a more local specific cell population that the broad cell-type classifications.

#### Regressing Cell Cycle

All droplet-based datasets (**Table 2**) were subject to cell-cycle regression using both Seurat and CycleMix. For Seurat, we used sctransform including the phase scores for S and G2M in the model as the creators recommend. For CycleMix, we specifically regressed differences between S and G2M cells while preserving differences between cycling and non-cycling cells (see above). Seurat requires datasets to be randomly downsampled to no more than 5,000 cells for cell-cycle regression, thus, we did the same when running the CycleMix regression to ensure consistency. For CycleMix, we downsampled to ensure equal presentation of each cell-cycle phase. Fit coefficients were used to regress the cellcycle from all cells. We then assessed the purity of neighbourhoods around G2M cells using the method described above. We assessed both purity of G2M cells and purity of either G2M or S cells to assess whether differences between cycling and non-cycling cells were preserved by each method.

For each dataset we compared the inferred frequency of cycling cells (S/G2M) from each method between proliferative and quiescent cell-types using a two-sided proportion test. Cell-types with fewer than 50 cells were excluded from analysis. Proliferation scores were calculated for each cell based on the highest phase score from either S or G2M and distributions of proliferative scores were compared using a KS-test.

#### Quiescence

Marker genes of quiescence were obtained from Cheung and Rando [40], and simplified to up (weight = 1), and down (weight = −1) and added to the Tirosh dataset of cell-cycle genes. CycleMix was used to classify each cell using this combined set of marker genes. Mouse orthologs of the Cheung genes were used for mouse datasets. The top quiescent cell-types were identified as those with the highest proportion of quiescent cells, excluding all cell-types with fewer than 50 cells.

Novel quiescence-associated genes were identified using a general linear model applied to log2 normalized expression values, including both sample ID and cell-type ID as covariates. Consensus genes were defined as those significantly up or down regulated in quiescent cells in 4/6 datasets. Pathway enrichments were determined using Fisher’s exact test on GO biological processes, Reactome, MSigdb Hallmark and Biocarta pathways.

#### Automatic Annotation

For each droplet-based dataset, the provided cell-type annotations were used to identify the top 10 up and top 10 down regulated genes in each cell type according to the difference in the proportion of cells expressing that gene. Each cell-type was downsampled randomly to 500 cells to define the marker genes. These 20 genes were assigned weights of +1/−1 and used to classify cells using CycleMix. Performance was measured using the adjusted rand (ARI) between classified cell-types and the author’s original annotations after excluding cells that were not classified to any cell-type.

#### iPSC-derived macrophages

Single-cell RNAseq data from iPSC-derived macrophages was obtained from the Gene Expression Omnibus (GSE250618). Data was filtered and normalized as previously described [49]. Seurat was run using the developer’s provided cell-cycle marker genes. CycleMix was run using the Tirosh cell-cycle geneset augmented with the G0 marker genes from Cheung et al. [40].

## Data Availability

List of accession numbers for benchmarking datasets are available in Table 1, real 10X datasets used for benchmarking (Table 2) are available in cellxgene at the provided links, and iPSC-derived macrophage data is available in Gene Expression Omnibus (GSE250618).

## Code Availability

CycleMix is freely available on Github under a GPLv3 open-source license: https://github.com/tallulandrews/CycleMix and is currently being submitted to Bioconductor.

## Author Contributions

TA conceptualized the project, designed analyses, performed analyses, and wrote the manuscript. JP designed analyses and performed analyses.

---

**Figure S1:**
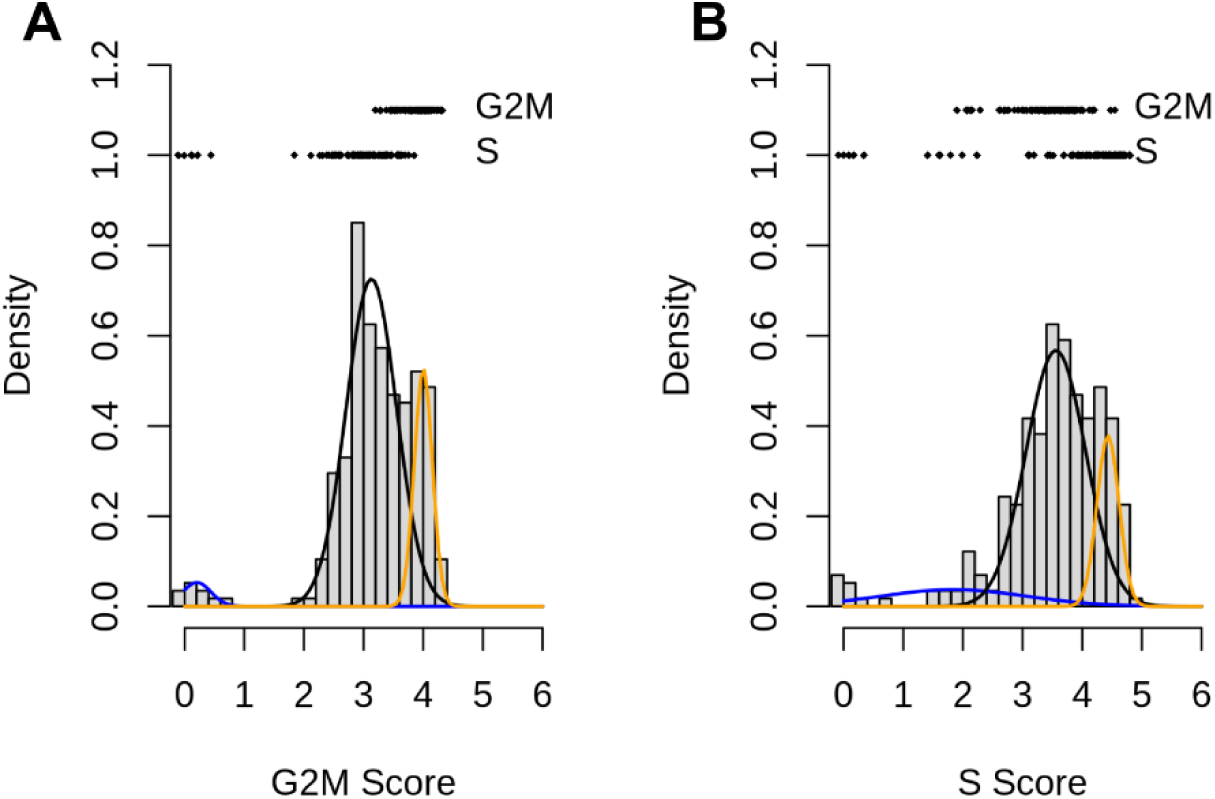
Buettner mESC cells classified into cell-cycle phases by CycleMix. (A) Optimal mixture model for G2M-phase score. (B) Optimal mixture model for S-phase score. Dots indicate the ground truth cell-cycle phase of each cell.

**Figure S2:**
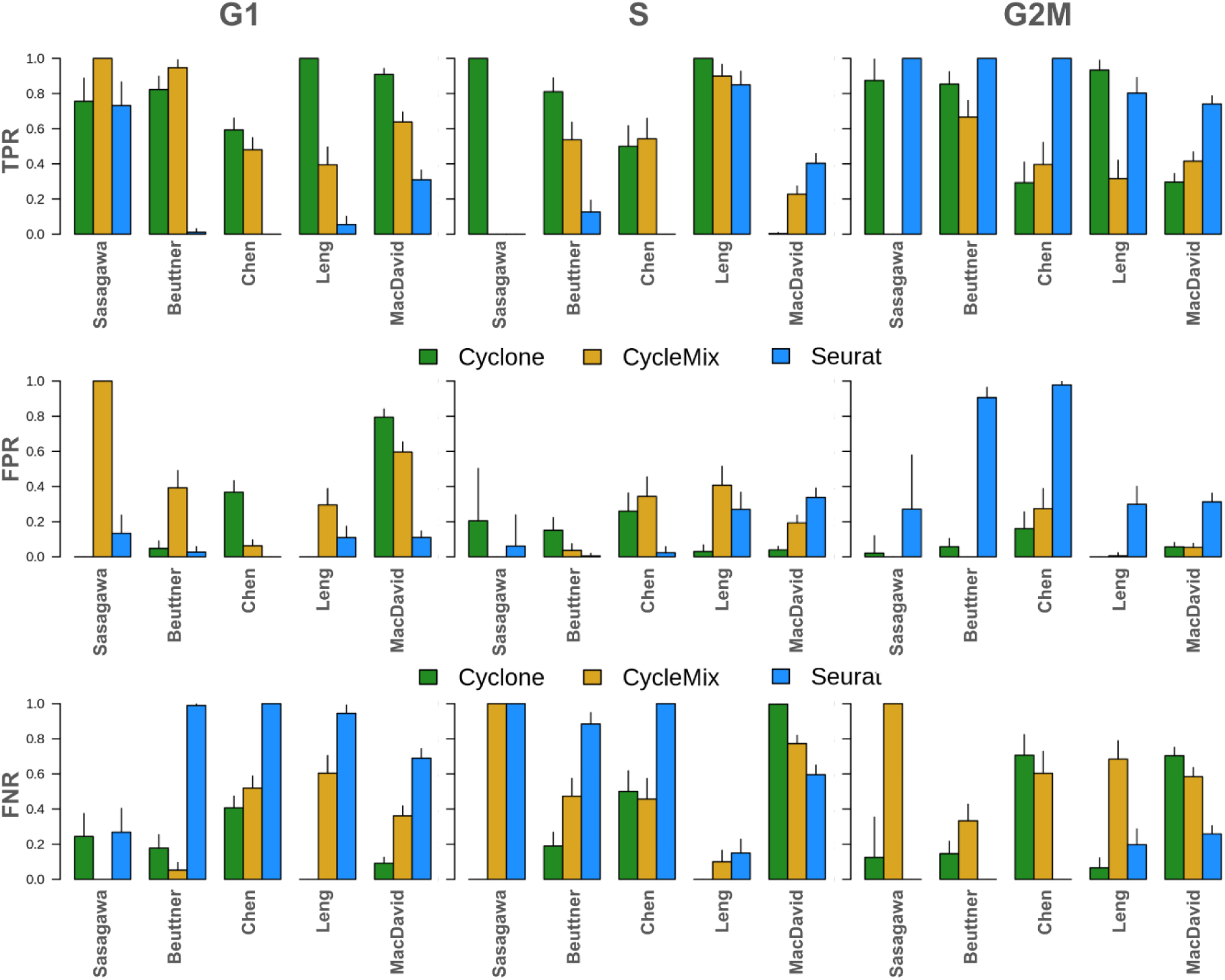
Accuracy of cell classifications on gold-standard ground truth datasets. (top) true positive rate (middle) false positive rate (bottom) false negative rate of each cell-cycle stage G1 (left), S (middle), and G2M (right)

**Figure S3:**
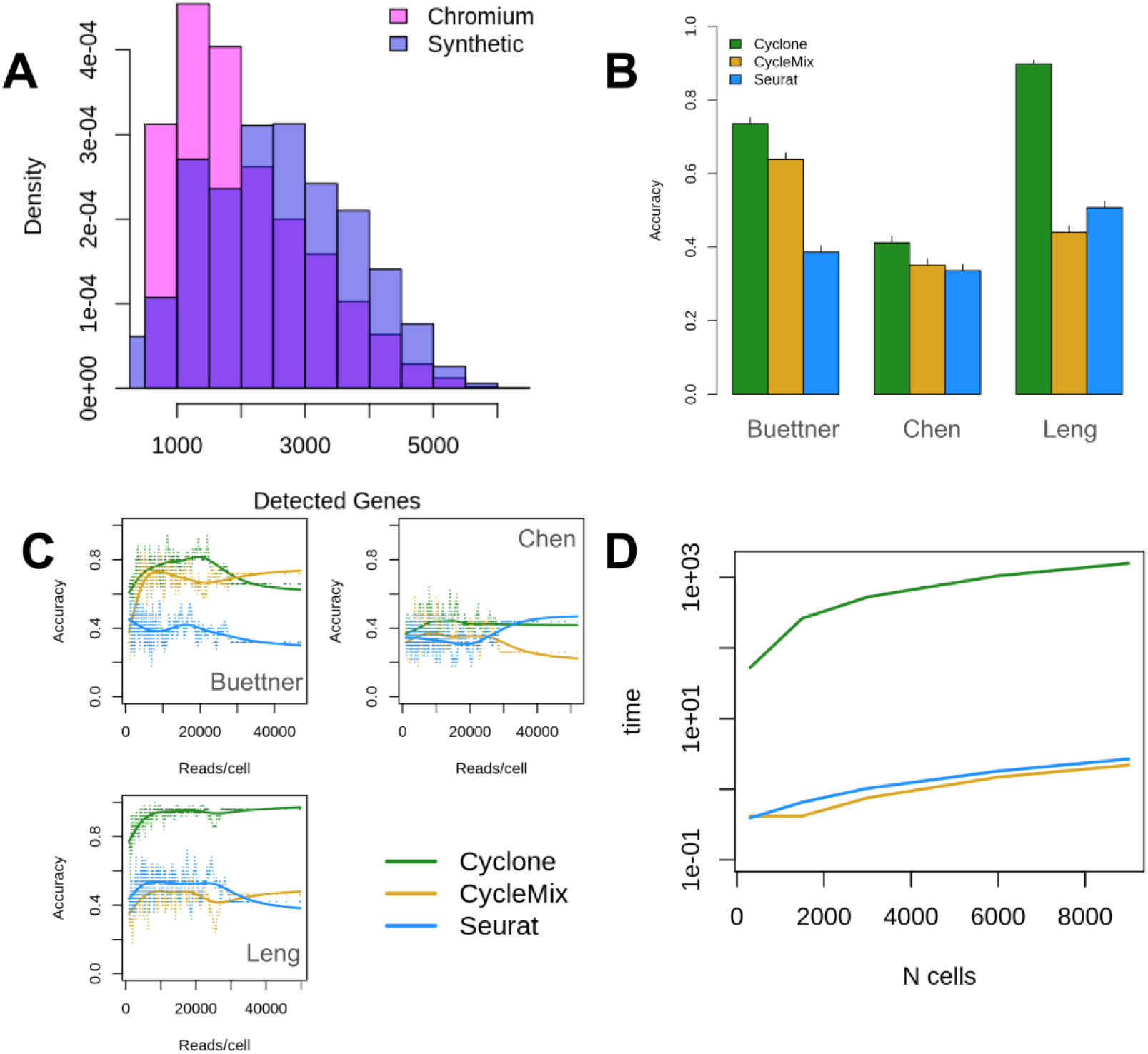
Benchmarking on resampled datasets to mimic droplet-based methods. (A) Downsampling of real datasets resulted in similar total detected genes per cell as 10X Chromium. (B) Cyclone generally outperformed other methods on all three downsampled datasets. Bars indicate 95% CI. (C) Relationship between classification accuracy and total reads / cell. Each point is average accuracy over 50 cells, and lines are spline fits to the trend across cells. (D) Cyclone was >500x slower than CycleMix or Seurat, requiring 30 minutes per 10,000 cells vs < 5 seconds per 10,000 cells for Seurat or CycleMix.

**Figure S4:**
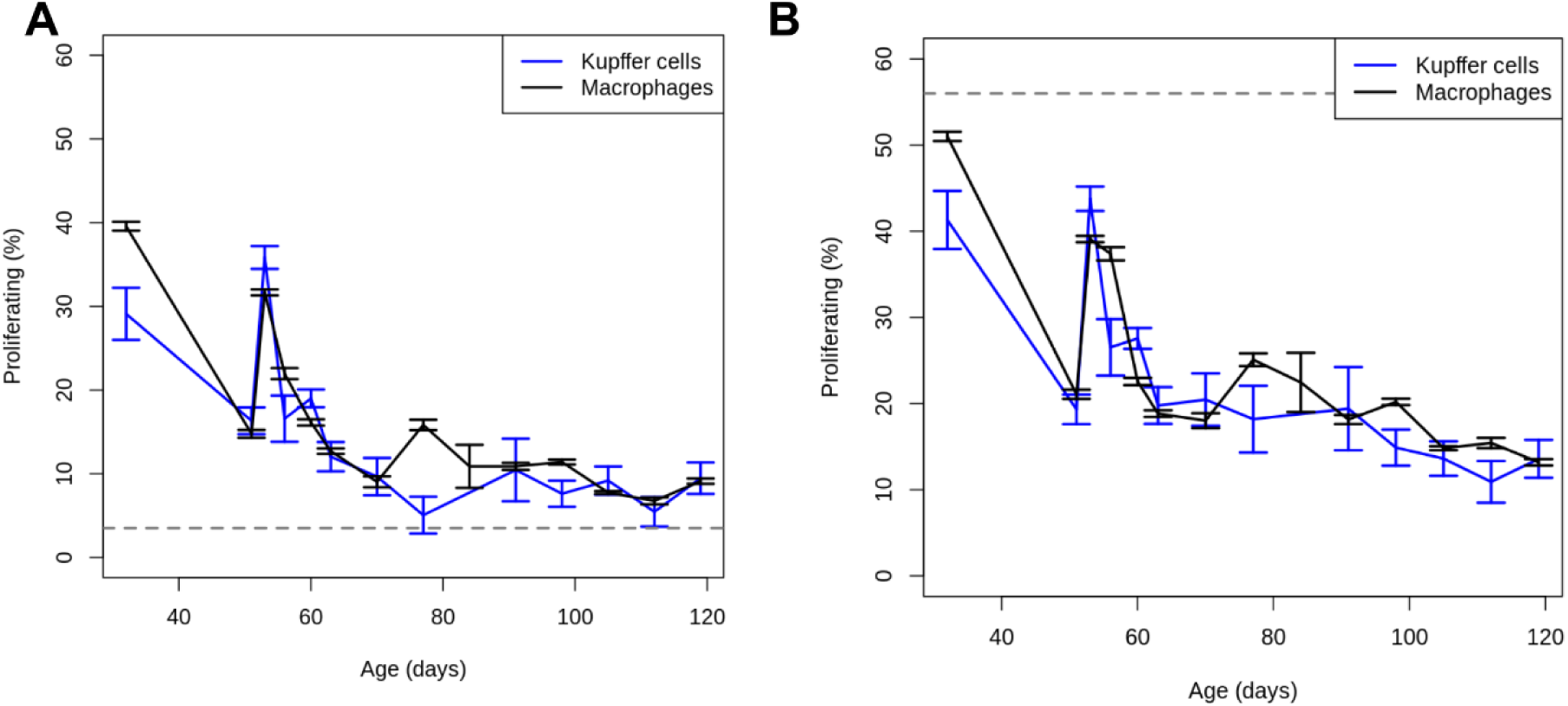
Proportion of fetal myeloid cells that are proliferating according to CycleMix & Seurat. (A) Proportion of proliferating macrophage or Kupffer cells across human fetal development using CycleMix. Dashed line indicates proportion of iPSC-derived Kupffer cells were proliferating. Error bars indicate 95% confidence intervals. (B) the same as A only using Seurat’s cell cycle classification instead.

**Figure S5:**
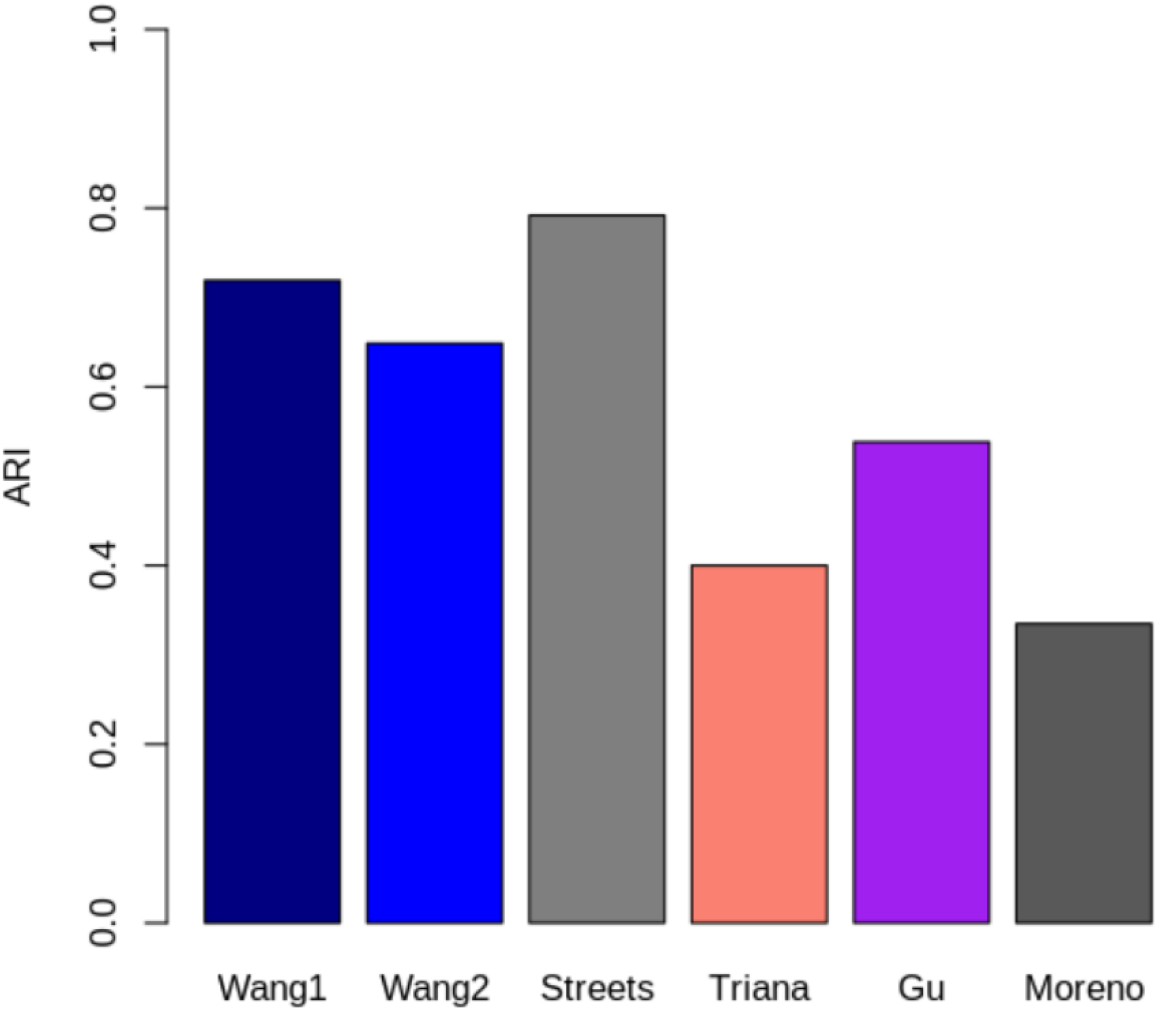
Accuracy of using CycleMix for cell-type annotation. Adjusted Rand Index (ARI) comparing CycleMix cell-type annotations to the original author annotations. Wang1 = colon, Wang2 = ileum.

## References

1. Fares J, Fares MY, Khachfe HH, Salhab HA, Fares Y. Molecular principles of metastasis: a hallmark of cancer revisited. Signal Transduct Target Ther. 2020;5: 28. doi:10.1038/s41392-020-0134-x

2. Hanahan D, Weinberg RA. Hallmarks of cancer: the next generation. Cell. 2011;144: 646–674. doi:10.1016/j.cell.2011.02.013

3. Joseph C, Mangani AS, Gupta V, Chitranshi N, Shen T, Dheer Y, et al. Cell cycle deficits in neurodegenerative disorders: uncovering molecular mechanisms to drive innovative therapeutic development. Aging Dis. 2020;11: 946–966. doi:10.14336/AD.2019.0923

4. Wang W, Bu B, Xie M, Zhang M, Yu Z, Tao D. Neural cell cycle dysregulation and central nervous system diseases. Prog Neurobiol. 2009;89: 1–17. doi:10.1016/j.pneurobio.2009.01.007

5. Lee K, Gusella GL, He JC. Epithelial proliferation and cell cycle dysregulation in kidney injury and disease. Kidney Int. 2021;100: 67–78. doi:10.1016/j.kint.2021.03.024

6. Katsel P, Davis KL, Li C, Tan W, Greenstein E, Kleiner Hoffman LB, et al. Abnormal indices of cell cycle activity in schizophrenia and their potential association with oligodendrocytes. Neuropsychopharmacology. 2008;33: 2993–3009. doi:10.1038/npp.2008.19

7. Abou Chakra M, Isserlin R, Tran TN, Bader GD. Control of tissue development and cell diversity by cell cycle-dependent transcriptional filtering. eLife. 2021;10. doi:10.7554/eLife.64951

8. Kolodziejczyk AA, Kim JK, Tsang JCH, Ilicic T, Henriksson J, Natarajan KN, et al. Single Cell RNA-Sequencing of Pluripotent States Unlocks Modular Transcriptional Variation. Cell Stem Cell. 2015;17: 471–485. doi:10.1016/j.stem.2015.09.011

9. Blasi T, Buettner F, Strasser MK, Marr C, Theis FJ. cgCorrect: a method to correct for confounding cell-cell variation due to cell growth in single-cell transcriptomics. Phys Biol. 2017;14: 036001. doi:10.1088/1478-3975/aa609a

10. Szalai B, Subramanian V, Holland CH, Alföldi R, Puskás LG, Saez-Rodriguez J. Signatures of cell death and proliferation in perturbation transcriptomics data-from confounding factor to effective prediction. Nucleic Acids Res. 2019;47: 10010–10026. doi:10.1093/nar/gkz805

11. Andrews TS, Kiselev VY, McCarthy D, Hemberg M. Tutorial: guidelines for the computational analysis of single-cell RNA sequencing data. Nat Protoc. 2020;16: 1–9. doi:10.1038/s41596-020-00409-w

12. Kiselev VY, Andrews TS, Hemberg M. Challenges in unsupervised clustering of single-cell RNA-seq data. Nat Rev Genet. 2019;20: 273–282. doi:10.1038/s41576-018-0088-9

13. Heumos L, Schaar AC, Lance C, Litinetskaya A, Drost F, Zappia L, et al. Best practices for single-cell analysis across modalities. Nat Rev Genet. 2023;24: 550–572. doi:10.1038/s41576-023-00586-w

14. Luecken MD, Theis FJ. Current best practices in single-cell RNA-seq analysis: a tutorial. Mol Syst Biol. 2019;15: e8746. doi:10.15252/msb.20188746

15. Gulati GS, D’Silva JP, Liu Y, Wang L, Newman AM. Profiling cell identity and tissue architecture with single-cell and spatial transcriptomics. Nat Rev Mol Cell Biol. 2025;26: 11–31. doi:10.1038/s41580-024-00768-2

16. Andrews TS, Nakib D, Perciani CT, Ma XZ, Liu L, Winter E, et al. Single-cell, single-nucleus, and spatial transcriptomics characterization of the immunological landscape in the healthy and PSC human liver. J Hepatol. 2024;80: 730–743. doi:10.1016/j.jhep.2023.12.023

17. Almeida MC, Eger SJ, He C, Audouard M, Nikitina A, Glasauer SMK, et al. Single-nucleus RNA sequencing demonstrates an autosomal dominant Alzheimer’s disease profile and possible mechanisms of disease protection. Neuron. 2024. doi:10.1016/j.neuron.2024.02.009

18. Cui A, Huang T, Li S, Ma A, Pérez JL, Sander C, et al. Dictionary of immune responses to cytokines at single-cell resolution. Nature. 2024;625: 377–384. doi:10.1038/s41586-023-06816-9

19. Sikkema L, Strobl DC, Zappia L, Madissoon E, Markov NS, Zaragosi L-E, et al. An integrated cell atlas of the human lung in health and disease. BioRxiv. 2022. doi:10.1101/2022.03.10.483747

20. Zeisel A, Hochgerner H, Lönnerberg P, Johnsson A, Memic F, van der Zwan J, et al. Molecular architecture of the mouse nervous system. Cell. 2018;174: 999–1014.e22. doi:10.1016/j.cell.2018.06.021

21. McDavid A, Finak G, Gottardo R. The contribution of cell cycle to heterogeneity in single-cell RNA-seq data. Nat Biotechnol. 2016;34: 591–593. doi:10.1038/nbt.3498

22. Lauridsen FKB, Jensen TL, Rapin N, Aslan D, Wilhelmson AS, Pundhir S, et al. Differences in cell cycle status underlie transcriptional heterogeneity in the HSC compartment. Cell Rep. 2018;24: 766–780. doi:10.1016/j.celrep.2018.06.057

23. Gu Y, Bartolomé-Casado R, Xu C, Bertocchi A, Janney A, Heuberger C, et al. Immune microniches shape intestinal Treg function. Nature. 2024;628: 854–862. doi:10.1038/s41586-024-07251-0

24. Lönnberg T, Svensson V, James KR, Fernandez-Ruiz D, Sebina I, Montandon R, et al. Single-cell RNA-seq and computational analysis using temporal mixture modelling resolves Th1/Tfh fate bifurcation in malaria. Sci Immunol. 2017;2. doi:10.1126/sciimmunol.aal2192

25. Pu Y, Li L, Peng H, Liu L, Heymann D, Robert C, et al. Drug-tolerant persister cells in cancer: the cutting edges and future directions. Nat Rev Clin Oncol. 2023;20: 799–813. doi:10.1038/s41571-023-00815-5

26. Deshpande A, Sicinski P, Hinds PW. Cyclins and cdks in development and cancer: a perspective. Oncogene. 2005;24: 2909–2915. doi:10.1038/sj.onc.1208618

27. Sasagawa Y, Nikaido I, Hayashi T, Danno H, Uno KD, Imai T, et al. Quartz-Seq: a highly reproducible and sensitive single-cell RNA sequencing method, reveals non-genetic gene-expression heterogeneity. Genome Biol. 2013;14: R31. doi:10.1186/gb-2013-14-4-r31

28. Buettner F, Natarajan KN, Casale FP, Proserpio V, Scialdone A, Theis FJ, et al. Computational analysis of cell-to-cell heterogeneity in single-cell RNA-sequencing data reveals hidden subpopulations of cells. Nat Biotechnol. 2015;33: 155–160. doi:10.1038/nbt.3102

29. Leng N, Chu L-F, Barry C, Li Y, Choi J, Li X, et al. Oscope identifies oscillatory genes in unsynchronized single-cell RNA-seq experiments. Nat Methods. 2015;12: 947–950. doi:10.1038/nmeth.3549

30. McDavid A, Dennis L, Danaher P, Finak G, Krouse M, Wang A, et al. Modeling bi-modality improves characterization of cell cycle on gene expression in single cells. PLoS Comput Biol. 2014;10: e1003696. doi:10.1371/journal.pcbi.1003696

31. Riba A, Oravecz A, Durik M, Jiménez S, Alunni V, Cerciat M, et al. Cell cycle gene regulation dynamics revealed by RNA velocity and deep-learning. Nat Commun. 2022;13: 2865. doi:10.1038/s41467-022-30545-8

32. Scialdone A, Natarajan KN, Saraiva LR, Proserpio V, Teichmann SA, Stegle O, et al. Computational assignment of cell-cycle stage from single-cell transcriptome data. Methods. 2015;85: 54–61. doi:10.1016/j.ymeth.2015.06.021

33. Butler A, Hoffman P, Smibert P, Papalexi E, Satija R. Integrating single-cell transcriptomic data across different conditions, technologies, and species. Nat Biotechnol. 2018;36: 411–420. doi:10.1038/nbt.4096

34. Tirosh I, Venteicher AS, Hebert C, Escalante LE, Patel AP, Yizhak K, et al. Single-cell RNA-seq supports a developmental hierarchy in human oligodendroglioma. Nature. 2016;539: 309–313. doi:10.1038/nature20123

35. Macosko EZ, Basu A, Satija R, Nemesh J, Shekhar K, Goldman M, et al. Highly Parallel Genome-wide Expression Profiling of Individual Cells Using Nanoliter Droplets. Cell. 2015;161: 1202–1214. doi:10.1016/j.cell.2015.05.002

36. Satija R, Farrell JA, Gennert D, Schier AF, Regev A. Spatial reconstruction of single-cell gene expression data. Nat Biotechnol. 2015;33: 495–502. doi:10.1038/nbt.3192

37. Whitfield ML, Sherlock G, Saldanha AJ, Murray JI, Ball CA, Alexander KE, et al. Identification of genes periodically expressed in the human cell cycle and their expression in tumors. Mol Biol Cell. 2002;13: 1977–2000. doi:10.1091/mbc.02-02-0030.

38. Chervov A, Zinovyev A. Computational challenges of cell cycle analysis using single cell transcriptomics. arXiv. 2022. doi:10.48550/arxiv.2208.05229

39. Chen Y, Zheng Y, Gao Y, Lin Z, Yang S, Wang T, et al. Single-cell RNA-seq uncovers dynamic processes and critical regulators in mouse spermatogenesis. Cell Res. 2018;28: 879–896. doi:10.1038/s41422-018-0074-y

40. Cheung TH, Rando TA. Molecular regulation of stem cell quiescence. Nat Rev Mol Cell Biol. 2013;14: 329–340. doi:10.1038/nrm3591

41. Finak G, McDavid A, Yajima M, Deng J, Gersuk V, Shalek AK, et al. MAST: a flexible statistical framework for assessing transcriptional changes and characterizing heterogeneity in single-cell RNA sequencing data. Genome Biol. 2015;16: 278. doi:10.1186/s13059-015-0844-5

42. Clarke ZA, Bader G. MALAT1 expression indicates cell quality in single-cell RNA sequencing data. BioRxiv. 2024. doi:10.1101/2024.07.14.603469

43. Dhanasekaran R, Deutzmann A, Mahauad-Fernandez WD, Hansen AS, Gouw AM, Felsher DW. The MYC oncogene - the grand orchestrator of cancer growth and immune evasion. Nat Rev Clin Oncol. 2022;19: 23–36. doi:10.1038/s41571-021-00549-2

44. Dang CV. c-Myc target genes involved in cell growth, apoptosis, and metabolism. Mol Cell Biol. 1999;19: 1–11. doi:10.1128/MCB.19.1.1

45. Matthews HK, Bertoli C, de Bruin RAM. Cell cycle control in cancer. Nat Rev Mol Cell Biol. 2022;23: 74–88. doi:10.1038/s41580-021-00404-3

46. Ren B, Cam H, Takahashi Y, Volkert T, Terragni J, Young RA, et al. E2F integrates cell cycle progression with DNA repair, replication, and G(2)/M checkpoints. Genes Dev. 2002;16: 245–256. doi:10.1101/gad.949802

47. Gomez Perdiguero E, Klapproth K, Schulz C, Busch K, Azzoni E, Crozet L, et al. Tissue-resident macrophages originate from yolk-sac-derived erythro-myeloid progenitors. Nature. 2015;518: 547–551. doi:10.1038/nature13989

48. Mass E, Nimmerjahn F, Kierdorf K, Schlitzer A. Tissue-specific macrophages: how they develop and choreograph tissue biology. Nat Rev Immunol. 2023;23: 563–579. doi:10.1038/s41577-023-00848-y

49. Kent GM, Atkins MH, Lung B, Nikitina A, Fernandes IM, Kwan JJ, et al. Human liver sinusoidal endothelial cells support the development of functional human pluripotent stem cell-derived Kupffer cells. Cell Rep. 2024;43: 114629. doi:10.1016/j.celrep.2024.114629

50. Suo C, Dann E, Goh I, Jardine L, Kleshchevnikov V, Park J-E, et al. Mapping the developing human immune system across organs. Science. 2022;376: eabo0510. doi:10.1126/science.abo0510

51. Jiang R, Sun T, Song D, Li JJ. Statistics or biology: the zero-inflation controversy about scRNA-seq data. Genome Biol. 2022;23: 31. doi:10.1186/s13059-022-02601-5

52. Svensson V. Droplet scRNA-seq is not zero-inflated. Nat Biotechnol. 2020;38: 147–150. doi:10.1038/s41587-019-0379-5

53. Korsunsky I, Millard N, Fan J, Slowikowski K, Zhang F, Wei K, et al. Fast, sensitive and accurate integration of single-cell data with Harmony. Nat Methods. 2019;16: 1289–1296. doi:10.1038/s41592-019-0619-0

54. Coronado D, Godet M, Bourillot P-Y, Tapponnier Y, Bernat A, Petit M, et al. A short G1 phase is an intrinsic determinant of naïve embryonic stem cell pluripotency. Stem Cell Res. 2013;10: 118–131. doi:10.1016/j.scr.2012.10.004

55. Becker KA, Ghule PN, Therrien JA, Lian JB, Stein JL, van Wijnen AJ, et al. Self-renewal of human embryonic stem cells is supported by a shortened G1 cell cycle phase. J Cell Physiol. 2006;209: 883–893. doi:10.1002/jcp.20776

56. Tirosh I, Izar B, Prakadan SM, Wadsworth MH, Treacy D, Trombetta JJ, et al. Dissecting the multicellular ecosystem of metastatic melanoma by single-cell RNA-seq. Science. 2016;352: 189–196. doi:10.1126/science.aad0501

57. Chen B, Wang J, Dai D, Zhou Q, Guo X, Tian Z, et al. AHNAK suppresses tumour proliferation and invasion by targeting multiple pathways in triple-negative breast cancer. J Exp Clin Cancer Res. 2017;36: 65. doi:10.1186/s13046-017-0522-4

58. Lee IH, Sohn M, Lim HJ, Yoon S, Oh H, Shin S, et al. Ahnak functions as a tumor suppressor via modulation of TGFβ/Smad signaling pathway. Oncogene. 2014;33: 4675–4684. doi:10.1038/onc.2014.69

59. Davis TA, Loos B, Engelbrecht AM. AHNAK: the giant jack of all trades. Cell Signal. 2014;26: 2683–2693. doi:10.1016/j.cellsig.2014.08.017

60. Smith LK, He Y, Park J-S, Bieri G, Snethlage CE, Lin K, et al. β2-microglobulin is a systemic pro-aging factor that impairs cognitive function and neurogenesis. Nat Med. 2015;21: 932–937. doi:10.1038/nm.3898

61. Clarke ZA, Andrews TS, Atif J, Pouyabahar D, Innes BT, MacParland SA, et al. Tutorial: guidelines for annotating single-cell transcriptomic maps using automated and manual methods. Nat Protoc. 2021;16: 2749–2764. doi:10.1038/s41596-021-00534-0

62. Li Z, Li F, Pan C, He Z, Pan X, Zhu Q, et al. Tumor cell proliferation (Ki-67) expression and its prognostic significance in histological subtypes of lung adenocarcinoma. Lung Cancer. 2021;154: 69–75. doi:10.1016/j.lungcan.2021.02.009

63. Nagy Á, Munkácsy G, Győrffy B. Pancancer survival analysis of cancer hallmark genes. Sci Rep. 2021;11: 6047. doi:10.1038/s41598-021-84787-5

64. Wang Z, Zhang W, Chen L, Lu X, Tu Y. Lymphopenia in sepsis: a narrative review. Crit Care. 2024;28: 315. doi:10.1186/s13054-024-05099-4

65. Cheung Y, Jia H. A Unified Metric for Categorical and Numerical Attributes in Data Clustering. In: Pei J, Tseng VS, Cao L, Motoda H, Xu G, editors. Advances in Knowledge Discovery and Data Mining. Berlin, Heidelberg: Springer Berlin Heidelberg; 2013. pp. 135–146. doi:10.1007/978-3-642-37456-2_12

66. Wang Y, Song W, Wang J, Wang T, Xiong X, Qi Z, et al. Single-cell transcriptome analysis reveals differential nutrient absorption functions in human intestine. J Exp Med. 2020;217. doi:10.1084/jem.20191130

67. Megill C, Martin B, Weaver C, Bell S, Prins L, Badajoz S, et al. cellxgene: a performant, scalable exploration platform for high dimensional sparse matrices. BioRxiv. 2021. doi:10.1101/2021.04.05.438318

68. Ruiz Moreno C, Stunnenberg HG, Nilsson M, Brandner S, Kranendonk ME, Samuelsson E, et al. Harmonized single-cell landscape, intercellular crosstalk and tumor architecture of glioblastoma. BioRxiv. 2022. doi:10.1101/2022.08.27.505439

69. Gupta A, Efthymiou V, Kodani SD, Shamsi F, Patti ME, Tseng Y-H, et al. Mapping the transcriptional landscape of human white and brown adipogenesis using single-nuclei RNA-seq. Mol Metab. 2023;74: 101746. doi:10.1016/j.molmet.2023.101746

70. Triana S, Vonficht D, Jopp-Saile L, Raffel S, Lutz R, Leonce D, et al. Single-cell proteo-genomic reference maps of the hematopoietic system enable the purification and massive profiling of precisely defined cell states. Nat Immunol. 2021;22: 1577–1589. doi:10.1038/s41590-021-01059-0

